# Control of GSK3β nuclear localization by amino acid signaling requires GATOR1 but is mTORC1-independent

**DOI:** 10.1101/2024.04.09.588669

**Authors:** Diana Schwendener Forkel, Osemudiamen Ibazebo, Sumaiya Soha, Stephen J. Bautista, Stefania Impellizzeri, Roberto J. Botelho, Geoffrey Hesketh, Anne-Claude Gingras, Costin N. Antonescu

## Abstract

The availability of certain amino acids regulates cell survival, proliferation, growth, differentiation, and other cellular functions. Sensing of amino acids that converges on the GATOR1 and GATOR2 complexes supports activation of mTORC1 during amino acid replete conditions. Whether amino acid-derived cues regulate additional pathways remains poorly understood. We uncover that amino acid sensing involving GATOR1 and GATOR2 regulates the cellular localization of glycogen synthase kinase 3β (GSK3β). GATOR1 is required to recruit a subset of GSK3β to the lysosome selectively in the presence of amino acids. In addition, while under nutrient replete conditions GSK3β is largely cytosolic, amino acid starvation drives a portion of GSK3β into the nucleus. Acute replenishment of specific amino acids in starved cells triggered nuclear exit of GSK3β. This amino acid-stimulated GSKβ nuclear exit required GATOR1 and GATOR2 but was independent of mTORC1 and its activating RagA/B GTPases. This suggests that GATOR1 has a function that diverges from control of mTORC1 to regulate the nucleocytoplasmic shuttling of GSK3β. Furthermore, experimental restriction of GSK3β to the cytoplasm decreased cell survival in amino acid deficient conditions. This suggests that control of GSK3β nuclear localization by GATOR-dependent signals represents a cellular adaptation to metabolic stress that supports cell survival.

## Introduction

Cells can experience and must adapt to a variety of environmental conditions, including variations in the availability of nutrients such as amino acids ^1,2^. Nutrient sensing and response mechanisms regulate not only cell growth and survival during nutrient stress but also function as signals to control phenomena such as immune cell function ^3^ and cellular differentiation ^4^. In addition, nutrient sensing and adaptation to restricted nutrient availability in the tumor microenvironment support tumor progression in various forms of cancer ^5,6^.

A series of amino acid sensing complexes operate at the lysosome and converge on the activation of mechanistic target of rapamycin complex 1 (mTORC1) ^7^. Central among these is the GATOR1 complex ^8,9^, which contains the subunits DEPDC5, NPRL3, and NPRL2, the latter harboring GAP activity towards the GTPases RagA/B ^10^. Disruption of any of these subunits of GATOR1 leads to mTORC1 activation, indicating that all are required for normal function of GATOR1. In turn, RagA/B controls mTORC1 activation ^11,12^. Under conditions of amino acid scarcity, the GAP activity of GATOR1 promotes GTP hydrolysis by RagA/B, maintaining mTORC1 in the inactive state.

Specific amino acids such as leucine and arginine are sensed at the lysosome ^13,14^ by Sestrin2 and Castor, respectively ^15,16^. In the presence of leucine or arginine, Sestrin2 or Castor trigger the suppression of GATOR1 GAP activity, leading in turn to enhanced GTP binding of RagA/B. GTP-bound RagA/B lead to lysosomal recruitment of mTORC1, supporting mTORC1 activation^9^. Linking amino acid sensing by Castor and Sestrin2 to regulation of GATOR1 is the GATOR2 complex. The RING domains of three GATOR2 complex proteins Mios, WDR24, and WDR59 mediate amino acid regulation of GATOR1 that may involve ubiquitylation of the GATOR1 subunit NPRL2 ^17,18^. Thus, loss of GATOR2 complex proteins such as Mios suppresses mTORC1 activation in response to amino acids ^19^. Mitogenic signaling by phosphoinositide-3-kinase (PI3K) and Akt to activate Rheb is also required for mTORC1 activation ^20–22^.

This makes lysosomal sensing of amino acid availability and convergence of signals on GATOR1 a critical component in the regulation of cellular adaptation to amino acid availability. Interestingly, a recent study reported an mTORC1-independent role of DEPDC5 in GABA signaling and gene expression ^23^. Whether and how DEPDC5 may have additional mTORC1-independent roles in response to nutrient cues remains poorly understood.

Glycogen synthase kinase 3 β (GSK3β) is a serine/threonine kinase that can phosphorylate and regulate a range of substrates, many of which are implicated in regulation of cell metabolism ^24–27^. This includes phosphorylation of c-myc, Snail, C/EBPα and β, CREB, GTF2F1, and FOXK1 ^28–34^. GSK3β-dependent phosphorylation leads to degradation of c-myc ^31,35^ and this may allow GSK3β to regulate expression of glycolytic and other metabolic enzymes known to be controlled by c-myc ^36–38^. Interestingly, GSK3β negatively regulates one-carbon metabolism at least in part by suppressing the expression of several genes that regulate one-carbon metabolism ^39^. Collectively, these studies suggest that GSK3β may be important for metabolic adaptation, but the mechanisms that regulate GSK3β, allowing it to respond to nutrient signaling remain incompletely resolved.

Compared to other kinases, GSK3β has been reported to have an unusually high number of substrates ^24,27,40^, indicating that GSK3β should be subject to right regulation. Surprisingly, GSK3β is constitutively active, and while inhibitory phosphorylation on S9 by Akt and other kinases does suppress GSK3β activity ^41–46^, this inhibition is incomplete and may impact only a subset of GSK3β ^34^. Control of nucleocytoplasmic shuttling of GSK3β restricts access of GSK3β to specific substrates. Specifically, nuclear localization of GSK3β is required for phosphorylation and regulation of c-myc and other nuclear factors, while cytosolic GSK3β accesses other substrates. We and others reported that mTORC1 promotes exit of GSK3β into the cytosol ^33,34,39^ and that inhibition of mTORC1 or disruption of lysosomal traffic by expression of a mutant of Rab7 or loss of lysosomal acidification leads to nuclear translocation of GSK3β ^34^. Lysosomes are heterogeneous varying in size, pH, and cargo composition, and form a continuum along with late endosomes and their hybrid endo-lysosomes (reviewed by ^47^). Here, we use the term lysosome broadly to encompass the continuum of endo-lysosomes. These studies indicate that lysosomal signals control nucleocytoplasmic shuttling of GSK3β and that mTORC1 contributes to this regulation. However, it is not known if other signals derived from the lysosome may also contribute to regulation of GSK3β nucleocytoplasmic shuttling, in particular under conditions of amino acid starvation.

We thus examined the regulation of GSK3β nucleocytoplasmic shuttling by cues derived from amino acid availability. By tracking endogenous GSK3β in ARPE-19 epithelial cells and MDA-MB-231 breast cancer cells by immunofluorescence methods that we previously validated ^34^, we find that serum starved cells accumulated nuclear GSK3β, as we previously reported ^34^. Treatment of starved cells with only specific amino acids, leucine or arginine, triggered loss of GSK3β from the nucleus. Using pharmacological inhibition and siRNA gene silencing approaches, we dissect how mTORC1 and GATOR1/2 complexes control GSK3β nuclear localization in response to amino acid cues. We further examine how nucleocytoplasmic shuttling of GSK3β impacts proliferation and survival in the breast cancer cell line MDA-MB-231.

## Results

To examine how nutrient signals impact GSK3β nucleocytoplasmic localization, we examined GSK3β nuclear localization under various conditions of nutrient availability. Compared to cells in full DMEM/F12 media containing both serum and amino acids, MDA-MB-231 breast cancer cells incubated in EBSS minimal media for 1h exhibited a robust increase in nuclear GSK3β (**Figure 1A**). Notably, we have previously validated this strategy for antibody labeling of endogenous GSK3β ^34^. We also previously demonstrated that monitoring the nuclear localization of GSK3β by immunofluorescence by this strategy reported nuclear GSK3β levels, as this was sensitive to interference by expression of Ran GTPase mutants and supported by cell fractionation approaches ^34^. These current results are consistent with our previous observations that withdrawal of growth factor and amino acid signaling, which suppressed mTORC1, leads to robust nuclear GSK3β localization ^34^.

**Figure 1.**
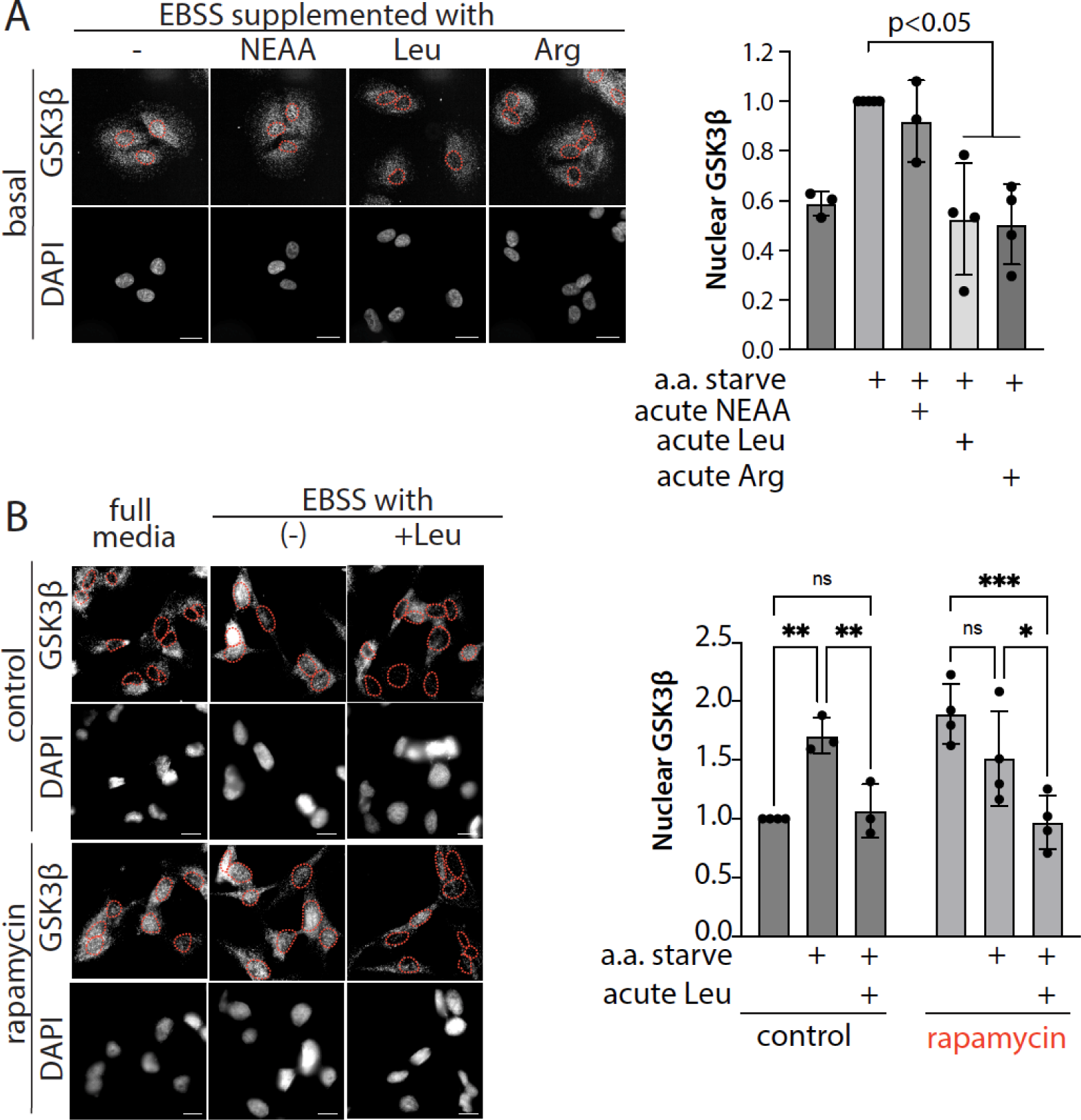
Amino acid availability controls GSK3β nuclear localization. (***A***) MDA-MB-231 cells were subjected to full media, amino acid starvation, or amino acid starvation followed by refeeding with leucine, arginine, or non-essential amino acids, as described in *Methods*. (***B***) MDA-MB-231 cells were treated as in (A), with some samples also being subject to treatment with 1 µM rapamycin for 1 h. For both (A) and (B), cells were fixed and labelled to detect endogenous GSK3β; shown (left panels) are representative micrographs obtained by widefield epifluorescence microscopy, scale 20 µm and (right panels) quantification of the mean fluorescence intensity of GSK3β in the nucleus depicting the mean ± SD from n = 3-4 independent experiments; each condition in each experiment determined by the median value from 30-40 individual cells. *, p < 0.05,

To next determine how GSK3β nuclear localization would be impacted by addition of specific amino acids to starved cells, we performed supplementation of the EBSS media with Leucine or Arginine, established ligands for Sestrin2 and Castor, regulators of GATOR complexes ^15,16^. We observed that either Leucine or Arginine supplementation re-localized GSK3β from the nucleus to the cytosol in EBSS-treated cells (**Figure 1A**). In contrast, supplementation with non-essential amino acids, which do not have known sensor(s) for signaling to the GATOR complexes, did not impact GSK3β nuclear localization in EBSS media (**Figure 1A**). These results indicate that GSK3β nuclear localization is acutely regulated by amino acids that signal through the GATOR complexes. However, whether and how this amino acid signaling to regulate GSK3β nuclear localization requires mTORC1 is not known.

### Regulation of GSK3β nuclear localization by amino acids is mTORC1-independent

To determine if the amino acid-induced re-localization of GSK3β from the nucleus to the cytosol by leucine is mTORC1-dependent, we examined the effect of treatment with the mTORC1 inhibitor rapamycin in MDA-MB-231 cells. Cells treated with 1h rapamycin alone or subjected to amino acid starvation for 1h and then subsequently treated with rapamycin for 1h exhibited a robust increase in nuclear GSK3β compared to cells in full media, as we reported previously ^34^ (**Figure 1B**). Importantly, cells that had been first starved of amino acids and treated with rapamycin exhibited loss of nuclear GSK3β upon acute (30 min) addition of leucine (**Figure 1B**). This indicates that in amino acid starved cells, acute replenishment of the essential amino acid leucine is sufficient to trigger signals that lead to complete return of nuclear GSK3β to the cytoplasm. Critically, the observation that the addition of leucine to amino acid starved cells elicits transit of nuclear GSK3β to the cytoplasm in cells also treated with rapamycin indicates that the signals that regulate GSK3β nucleocytoplasmic shuttling in amino acid starved cells by leucine addition are mTORC1-independent.

We observed similar effects of rapamycin on GSK3β nuclear exit triggered by leucine addition in ARPE-19 cells, a non-transformed cell line of epithelial lineage (**Figure S1**). We also found similar results using Torin-1 to inhibit mTORC1 instead of rapamycin (**Figure S1**). These results suggest that while mTORC1 controls GSK3β nucleocytoplasmic shuttling in cells in amino acid replete media ^34^, in cells subjected to amino acid starvation, an mTORC1-independent signaling pathway couples sensing of essential amino acids such as leucine and arginine to translocation of GSK3β from the nucleus to the cytoplasm.

### A subset of GSK3β is recruited to the lysosome in a GATOR1- and amino acid-dependent manner

We next sought to examine how lysosome-localized amino acid sensing signals control GSK3β localization, since we previously observed that a subset of GSK3β localized to the lysosome, and disruption of lysosomal traffic by Rab7 loss-of-function resulted in a robust increase in nuclear GSK3β ^34^. We focused on GATOR1 since this complex is localized to the lysosome and is regulated by leucine and arginine ^15,16^. We focused on ARPE-19 cells for these experiments since these cells are more spread out than MDA-MB-231 cells and thus allow more facile resolution of individual intracellular membrane compartments. We used antibody labeling of endogenous GSK3β and the endolysosomal marker LAMP1 to examine the localization of a subset of GSK3β to the lysosome, as we had done previously. We observed that amino acid starvation triggered loss of GSK3β localizing with LAMP1 at the lysosome (**Figure 2 and S2**). Addition of leucine to amino acid starved cells in the presence of rapamycin treatment triggered a recovery of the recruitment of a subset of GSK3β to the lysosome (**Figure 2**). Importantly, silencing of the GATOR1 complex core subunit DEPDC5 resulted in loss of GSK3β from localization with LAMP1 in all conditions observed (**Figure 2**). These results suggest that a pool of GSK3β is recruited to the lysosome in a GATOR1-dependent manner, and that the recruitment of GSK3β to the lysosome is dependent on amino acid availability.

**Figure 2.**
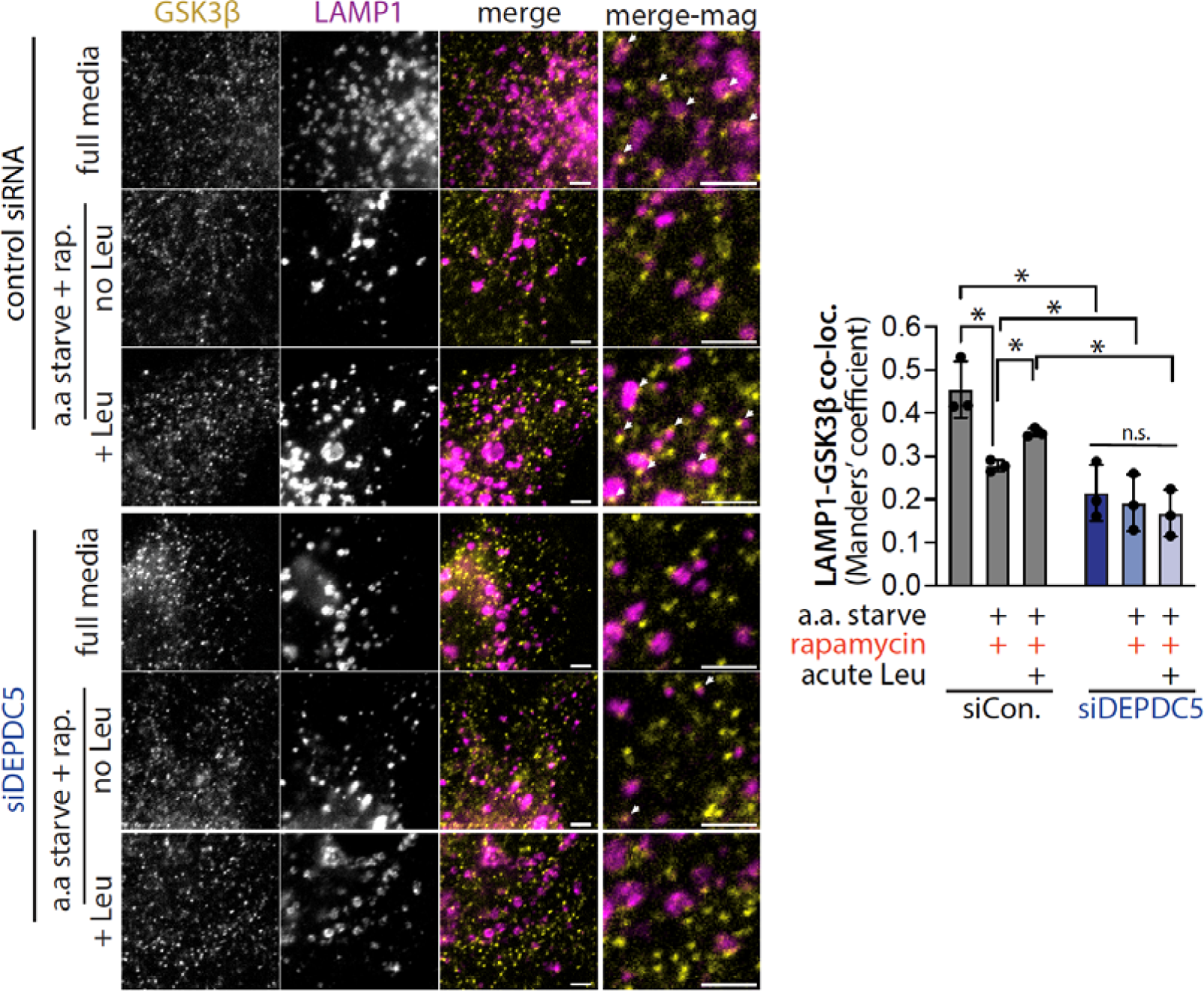
The GATOR1 subunit DEPDC5 is required for amino acid-dependent recruitment of a subset of GSK3β to the lysosome. ARPE-19 cells were transfected with siRNAs targeting DEPDC5 or non-targeting siRNA, and then subjected to full media, amino acid starvation, or amino acid starvation followed by refeeding with leucine, as described in *Methods*. Cells were fixed and labelled to detect endogenous GSK3β and LAMP1; shown (left panels) are representative micrographs obtained by widefield epifluorescence microscopy, Scale 5 µm and white arrowheads depicting manually-identified examples if overlap or proximity of GSK3β and LAMP1 signals and (right panels) quantification of Manders’ coefficient as the mean ± SD from n = 3 independent experiments; each condition in each experiment determined by the median value from 30-40 individual cells, *, p <0.05. See **Figure S2** for full images.

### Amino acid signaling does not appreciably alter DEPDC5 localization to the lysosome

These results indicate that a subset of GSK3β is localized to the lysosomes in a DEPDC5-dependent manner, and that this recruitment of GSK3β to lysosomes is suppressed by amino acid starvation. Some studies have reported that components of the GATOR1 complex such as NPRL2 and NPRL3 can localize to compartments other than the lysosome, including the nucleus, under some stress conditions ^48^. Since we observed that regulation of GSK3β localization at the lysosome was GATOR1-dependent, this suggests that it is possible that the loss of GSK3β from lysosomes upon amino acid starvation could reflect a loss of GATOR1 components including DEPDC5 from lysosomes. To examine this possibility, we used the Sleeping Beauty transposon system ^49^ to generate ARPE-19 cells that stably harbor a transgene for doxycycline-inducible DEPDC5-eGFP expression. To ensure non-perturbing levels of DEPDC5-eGFP expression, we used low doxycycline concentration to induce expression and used a non-perturbing method to enhance low fluorescence of green fluorochromes at the lysosomes by delivery of silver nanoparticles to the lysosome that we recently described ^50^, alongside labeling of lysosomes with Lysotracker Deep Red.

Using this approach and live-cell spinning disc confocal microscopy, we identified strong overlap of DEPDC5-eGFP with lysosomes as demarked by Lysotracker staining (**Figure 3 and S3**). Interestingly, we did not observe a drop in the overlap of DEPDC5-eGFP with lysosomes upon amino acid starvation or subsequent refeeding of starved cells with acute leucine supplementation. These results suggest that DEPDC5 regulates the recruitment of a subset of GSK3β to the lysosome, but that this does not result from loss of the localization of DEPDC5 itself to lysosomes upon conditions that trigger loss of GSK3β from these organelles. Instead, these results suggest that regulation of DEPDC5 conformation by amino acid signals could recruit a subset of GSK3β to the lysosome, which may in turn mediate regulation of broader GSK3β functions.

**Figure 3.**
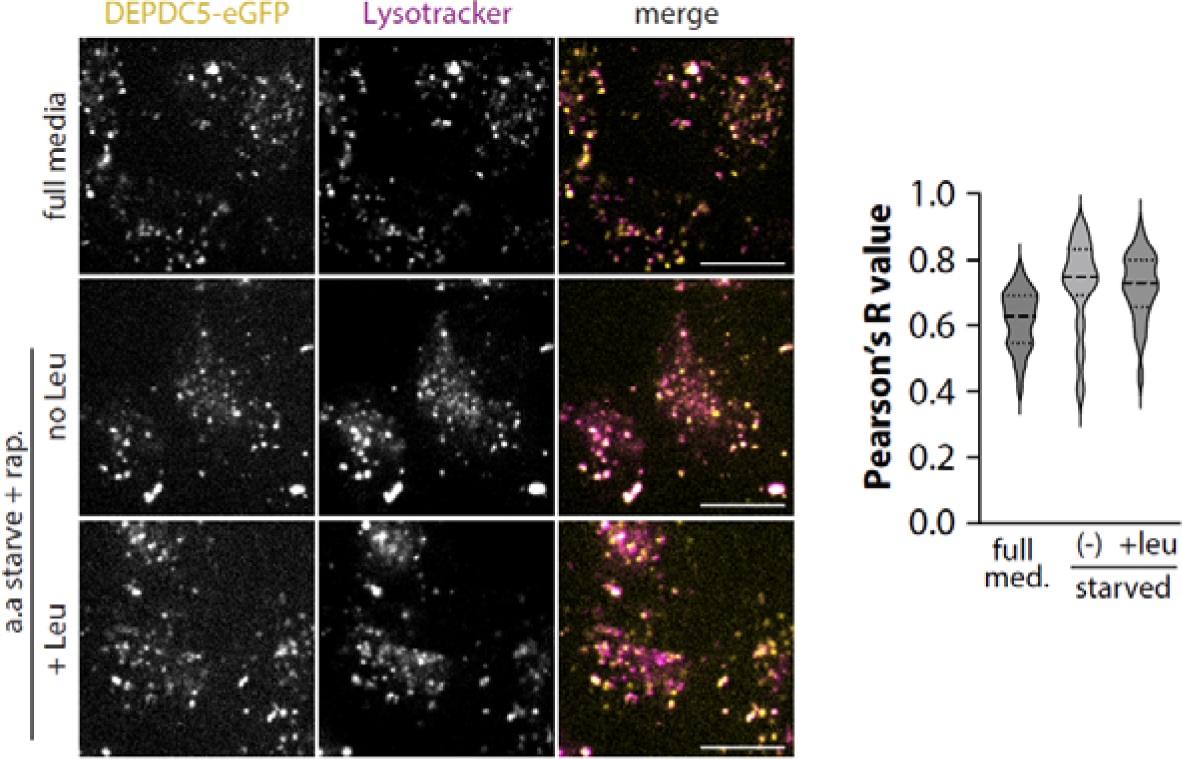
DEPDC5-eGFP localizes to lysosomes similarly under amino acid replete and starvation conditions. ARPE-19 eGFP-DEPDC5 cells were treated with 1 µg/mL doxycycline for 24 h to induce eGFP-DEPDC5 expression, treated with Lysotracker Deep Red, and then subjected to full media, amino acid starvation, or amino acid starvation followed by refeeding with leucine, as described in *Methods*. Shown (left panels) are representative micrographs obtained by widefield spinning-disc confocal microscopy, scale 20 µm and (right panels) quantification of colocalization in each condition by Pearson’s R coefficient. See **Figure S3** for full images.

### The regulation of GSK3β nuclear localization by amino acids is dependent on GATOR1 and GATOR2

The requirement for GATOR1 to allow a pool of GSK3β to be dynamically recruited to the surface of the lysosome in an amino-acid dependent manner suggests that GATOR1 may also be required for regulation of GSK3β nucleocytoplasmic shuttling by amino acid signaling. To examine this, we used silencing of DEPDC5 in ARPE-19 cells (**Figure S4**), and subjected cells to amino acid depletion, rapamycin treatment, and/or acute leucine addition as **in Figure 1**. DEPDC5 silencing was without effect on the levels of nuclear GSK3β in cells in full media or upon treatment with either rapamycin or amino acid starvation (**Figure 4A**). However, DEPDC5 siRNA silencing blocked the ability of acute leucine addition to amino acid starved and rapamycin treated cells to trigger nuclear exit of GSK3β, compared to control siRNA cells (**Figure 4A**). We observed similar results upon silencing of another GATOR1 complex subunit, NPRL2 (**Figure 4B**).

**Figure 4.**
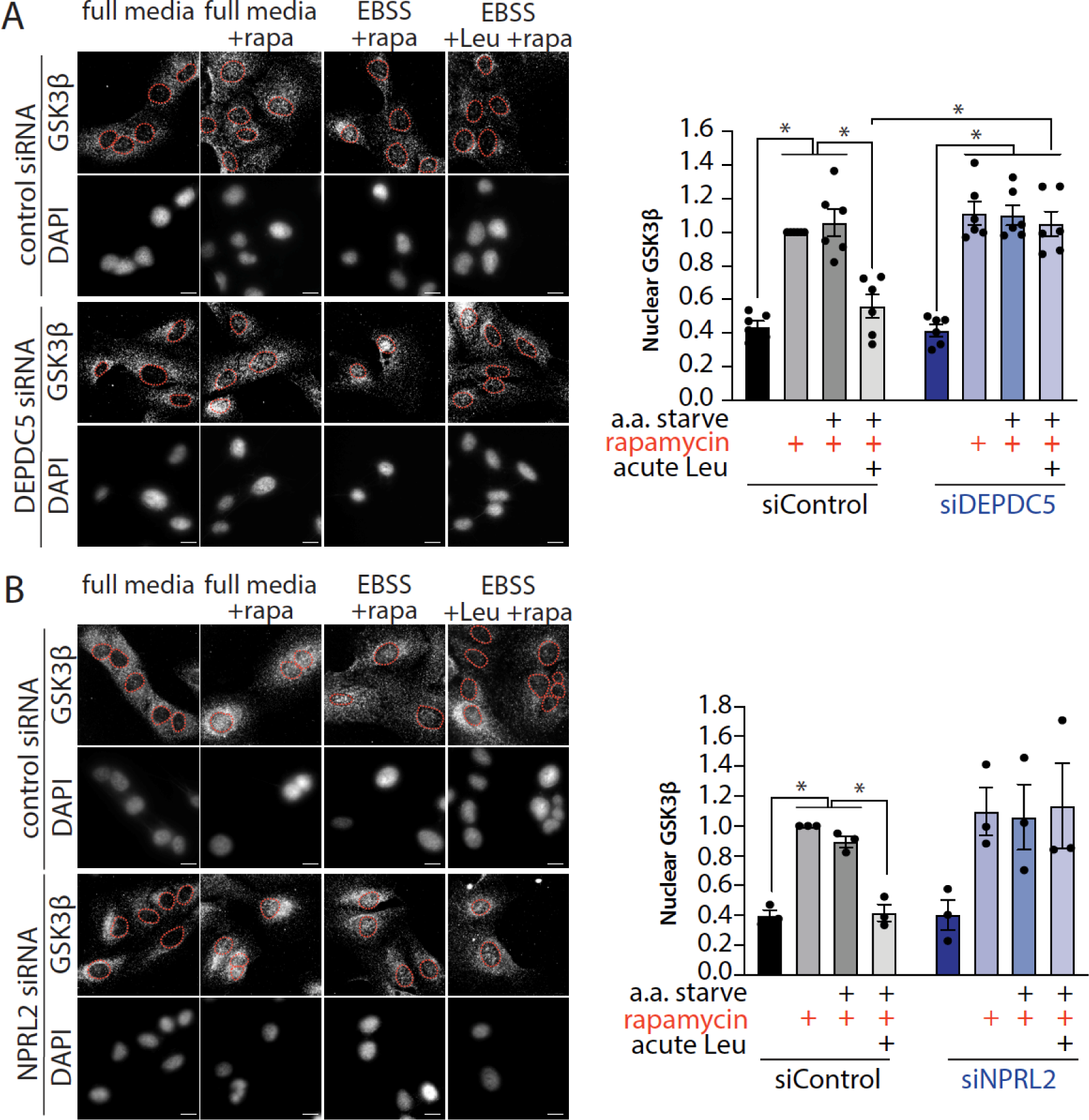
GATOR1 is required for amino acid-dependent control of GSK3β nuclear localization. ARPE-19 cells were transfected with siRNAs targeting DEPDC5 (***A***) or NPRL2 (***B***) or non-targeting siRNA, and then subjected to full media, amino acid starvation, or amino acid starvation followed by refeeding with leucine, as described in *Methods*. Cells were fixed and labelled to detect endogenous GSK3β; shown (left panels) are representative micrographs obtained by widefield epifluorescence microscopy, scale 20 µm and (right panels) quantification of the mean fluorescence intensity of GSK3β in the nucleus depicting the mean ± SD from n = 3-4 independent experiments; each condition in each experiment determined by the median value from 30-40 individual cells. *, p < 0.05.

Since leucine and arginine are sensed by Sestrin2 and Castor, respectively, thus in turn regulating GATOR2, we next examined the contribution of the core GATOR2 complex subunit Mios to amino acid-dependent regulation of GSK3β. Similar to the effect of silencing GATOR1 complex subunits, silencing Mios was without effect on the levels of nuclear GSK3β in cells in full media or upon treatment with either rapamycin or amino acid starvation (**Figure 5**). However, cells subjected to Mios silencing did not exhibit nuclear exit of GSK3β upon acute leucine treatment of amino acid starved, rapamycin-treated cells (**Figure 5**), an effect similar to silencing of GATOR1 components (**Figure 4**). These results indicate that amino acid sensing involving GATOR1 and GATOR2 is required to trigger nuclear exit of GSK3β in amino acid cells upon acute addition of amino acids, and that this occurs in an mTORC1-independent manner.

**Figure 5.**
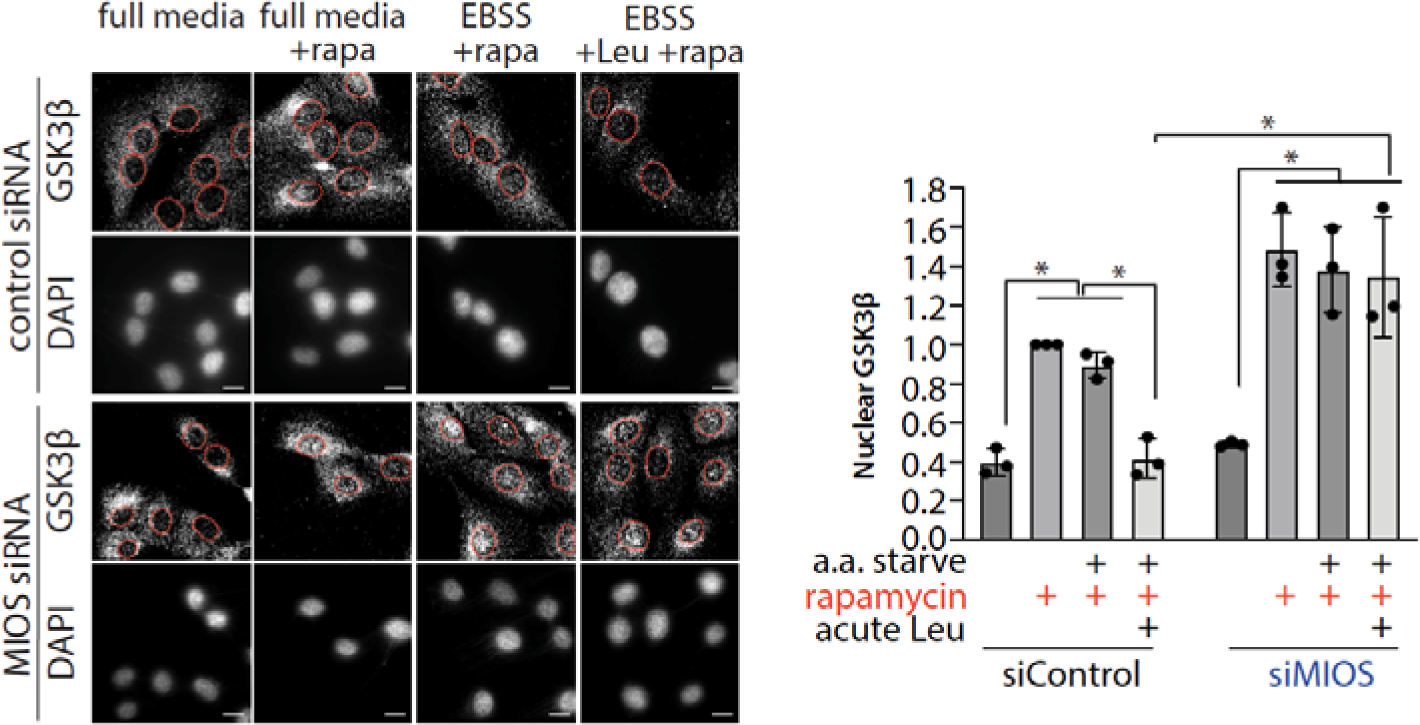
GATOR2 is required for amino acid-dependent control of GSK3β nuclear localization. ARPE-19 cells were transfected with siRNAs targeting Mios or non-targeting siRNA, and then subjected to full media, amino acid starvation, or amino acid starvation followed by refeeding with leucine, as described in *Methods*. Cells were fixed and labelled to detect endogenous GSK3β; shown (left panels) are representative micrographs obtained by widefield epifluorescence microscopy, scale 20 µm and (right panels) quantification of the mean fluorescence intensity of GSK3β in the nucleus depicting the mean ± SD from n = 3-4 independent experiments; each condition in each experiment determined by the median value from 30-40 individual cells. *, p < 0.05.

### The regulation of GSK3β nuclear localization by amino acids is independent of the RagA/B GTPases

Since the nuclear exit of GSK3β in amino acid starved cells by acute amino acid treatment is mTORC1-independent but requires GATOR1 and GATOR2, this raises the question of the role of Rag GTPases, as the latter are directly controlled by GATOR1 and in turn control mTORC1 activation. We subjected cells to silencing of RagA and RagB together, which was previously used to ascertain the involvement of Rag GTPases, for example in the lysosomal targeting of mTORC1 ^51^. Similar to silencing of GATOR1 and GATOR2 complex components, silencing of RagA/B was without effect on the levels of nuclear GSK3β in cells in full media or upon treatment with either rapamycin or amino acid starvation. However, and in contrast to the effects of silencing GATOR1 and GATOR2 complexes, silencing RagA/B was without effect on the nuclear exit of GSK3β upon acute leucine addition of amino acid starved, rapamycin-treated cells (**Figure 6**). These results are consistent with the nuclear exit of GSK3β by acute amino acid treatment being mTORC1-independent (**Figure 1**). This suggests that amino acid signaling bifurcates at GATOR1, with one arm leading to the regulation of Rag GTPases and mTORC1, and another arm exerting control over GSK3β nucleocytoplasmic shuttling in an mTORC1-independent manner.

**Figure 6.**
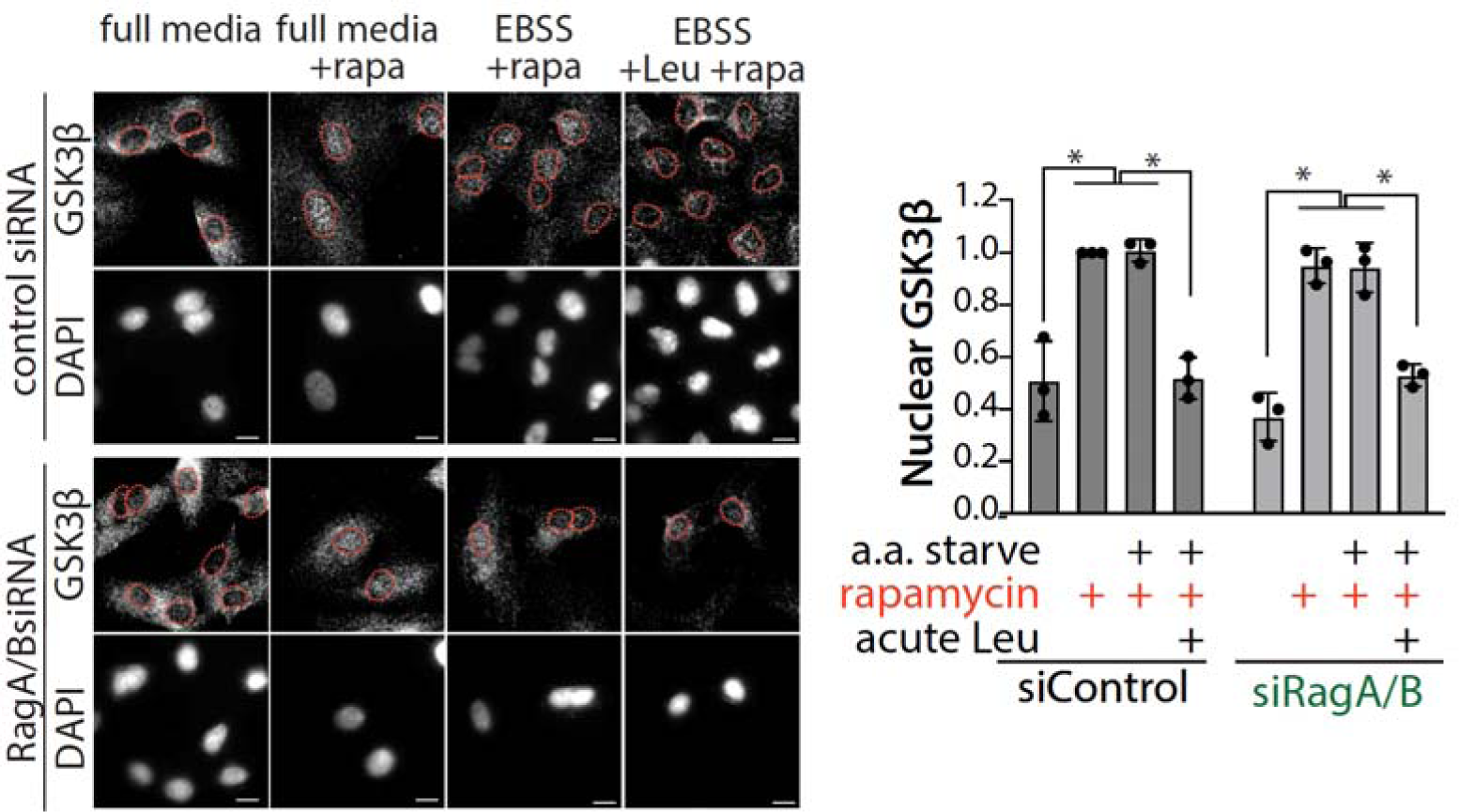
RagA/B are dispensable for amino acid-dependent control of GSK3β nuclear localization. ARPE-19 cells were transfected with siRNAs targeting both RagA and RagB or non-targeting siRNA, and then subjected to full media, amino acid starvation, or amino acid starvation followed by refeeding with leucine, as described in *Methods*. Cells were fixed and labelled to detect endogenous GSK3β; shown (left panels) are representative micrographs obtained by widefield epifluorescence microscopy, scale 20 µm and (right panels) quantification of the mean fluorescence intensity of GSK3β in the nucleus depicting the mean ± SD from n = 3-4 independent experiments; each condition in each experiment determined by the median value from 30-40 individual cells. *, p < 0.05.

### GSK3β nuclear localization controls cell proliferation and survival

Taken together, these results suggest that regulation of GSK3β nuclear localization is tightly regulated by amino acid signals. To determine how the regulation of GSK3β nuclear localization under various conditions of nutrient availability impacts cell physiology, we constructed a cell model that could uncouple GSK3β nuclear localization from amino acid sensing. We first used CRISPR/Cas9 to generate MDA-MB-231 cells with a GSK3β knockout (**Figure 7A**). Then, using this knockout cell model, we used the Sleeping Beauty transposon system to generate stable cells that allow doxycycline-inducible expression of GSK3β harboring nuclear localization sequence (GSK3β-NLS) or nuclear export sequence (GSK3β-NES). We confirmed that these cells exhibited GSK3β localization to the nucleus and cytoplasm, as expected (**Figure 7B**). We next subjected these cells to full media or amino acid and serum starvation conditions, henceforth referred to as “starvation conditions”. Of note, we supplemented the latter with 10 ng/mL EGF to provide some growth factor stimulation during these experiments that involved prolonged exposure to amino acid-free media.

**Figure 7.**
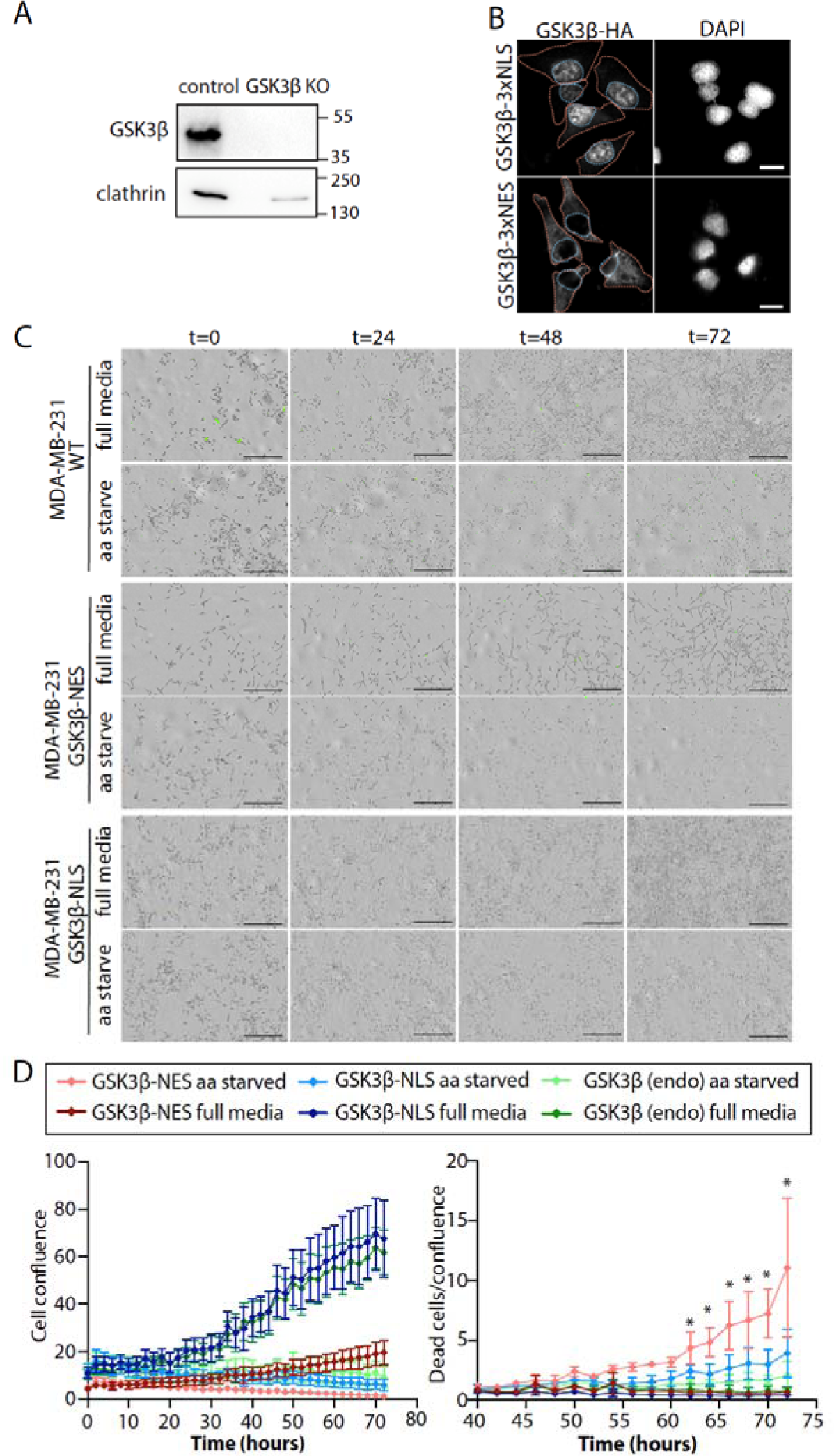
GSK3β with nuclear exclusion impairs MDA-MB-231 cell survival in amino acid starvation conditions. (***A***) MDA-MB-231 cells with GSK3β knockout were generated as described in Methods. Shown is a western blot detecting GSK3β expression. (***B***) MDA-MB-231 cells with GSK3β knockout were used to generate stable cells using the Sleeping Beauty transposing system for inducible expression of GSK3β with a nuclear localization sequence (GSK3β-NLS) or a nuclear export sequence (GSK3β-NES). Expression of each GSK3β construct was induced by 24h incubation in 250 ng/mL doxycycline, followed by immunofluorescence labeling using HA antibodies. Shown are representative micrographs obtained by widefield epifluorescence microscopy, Scale = 20 µm (***C-D***) MDA-MB-231 parental (wild-type) cells, or MDA-MB-231 with GSK3β knockout with rescue with GSK3β-NLS or GSK3β-NES induced with 250 ng/mL doxycycline were incubated in full media, amino acid starvation, or amino acid starvation supplemented with 10 ng/mL EGF. During this time, samples were also treated with CellTox green to visualize non-viable cells and subjected to imaging with an Incucyte SX5 microscope at 37C and 5% CO2 for 72 h. Shown in (C) are representative micrographs showing phase contrast images overlaid on CellTox green fluorescence signal. Scale 400 µm. Shown in (D, left panels) are the measurement of cell abundance determined by automated detection of cell confluence (left panels) and (D, right panels) non-viable cells relative to cell confluence; both as mean ± SD from 4 independent experiments. *, p < 0.05, (GSK3β-NES amino acid starved relative to all other treatments).

We subjected each condition to observation of cell confluence using automated tracing and cell viability using detection of labeling with CellTox Green (**Figure 7C**). As expected, each of the cell lines examined (MDA-MB-231 WT cells; GSK3β-NES cells; GSK3β-NLS cells) exhibited reduced cell abundance in starvation conditions compared with full media conditions. Interestingly, GSK3β-NES cells exhibited reduced cell confluence in both full media and starvation conditions compared to either the MDA-MB-231 WT cell line or the GSK3β-NLS cell line (**Figure 7D**, *left panels*). This suggests that the regulation of GSK3β nucleocytoplasmic shuttling regulates the proliferation and/or survival of MDA-MB-231 cells, as was previously reported ^33^. Notably, in amino acid starvation conditions, each cell line exhibited very little growth over the 72 h observation period, suggesting that cells arrested their cell cycle and/or that some conditions exhibited elevated levels of cell death.

To resolve whether these conditions triggered an increase in cell death in any of the conditions examined, we quantified the CellTox Green positive objects and normalized this to the cell confluence in each image (**Figure 7D**, *right panels*), as we have done previously ^52^. This allows the measurement of cell death indexed to a measure of total cell abundance. Strikingly, GSK3β-NLS grown in amino acid free media exhibited significantly more cell death than other conditions, including either GSK3β-NES cells or MDA-MB-231 WT cells grown in amino acid starvation media. This observation that the selective loss of cell viability in GSK3β-NES cells (that have cytosolic GSK3β) during amino acid starvation compared to cells with unperturbed GSK3β localization or cells with GSK3β restricted to the nucleus suggests that the translocation of GSK3β from the cytoplasm to the nucleus confers on cells an advantage under amino acid starvation.

## Discussion

We find that lysosomal amino acid sensing protein complexes GATOR1 and GATOR2 are required for the regulation of GSK3β nuclear localization in amino acid starved cells. We and others previously reported that GSK3β nucleocytoplasmic shuttling was controlled by mTORC1 under amino acid replete conditions ^33,34^. Specifically, treatment of various cells with the mTORC1 inhibitor rapamycin was sufficient to trigger strong accumulation of GSK3β in the nucleus, in amino acid replete conditions. In contrast, we now report that in amino acid starved cells, GSK3β nuclear exit triggered by acute treatment with leucine or arginine is insensitive to the mTORC1 inhibitors rapamycin (**Figure 1, S1**) or Torin1 (**Figure S1**). This suggests that two parallel mechanisms regulate GSK3β nuclear localization: (i) an mTORC1-dependent mechanism that operates in nutrient-replete conditions and (ii) an mTORC1-independent mechanism that relies on GATOR1 that operates in amino acid starved conditions.

### Mechanism of regulation of GSK3β by GATOR1/2 under amino acid starvation

We show that the GATOR1 subunit DEPDC5 is required for localization of a subset of GSK3β to the lysosome (**Figure 2**) under amino acid replete conditions. This indicates that GATOR1 may promote localization of GSK3β to the lysosome under nutrient replete conditions to support its regulation by mTORC1 specifically under this condition, although it does not appear that GSK3β is phosphorylated in an mTORC1-dependent manner ^34^. Hence, any regulation of GSK3β by mTORC1 in amino acid replete conditions is likely indirect and suggests the existence of a yet-to-be identified intermediate. Furthermore, we observed the mTORC1-independent and DEPDC5-dependent recruitment of a subset of GSK3β to the lysosome in amino acid starved cells upon acute leucine treatment (**Figure 2**). This suggests a regulatory mechanism at the level of GATOR1 that branches from regulation of Rag GTPases, which can direct GSK3β nuclear exit and lysosomal recruitment in a RagA/B- and mTORC1-independent manner. It is not clear if and how in amino acid starved cells, the leucine- or arginine-triggered recruitment of GSK3β to lysosomes may be related to the regulation GSK3β nucleocytoplasmic shuttling.

The previously described mTORC1-dependent ^33,34^ and the mTORC1-independent mechanism of regulation of GSK3β by amino acid signals identified in this study may each impact distinct stages of GSK3β nucleocytoplasmic shuttling. GSK3β has a nuclear localization sequence that is comprised of basic amino acids (85-103) ^53^; such basic nuclear localization sequences are recognized by importin α/β (Kapβ1 pathway) ^54^, although GSK3β may also binds to a different nuclear import receptor, Kapβ2 *via* a separate 109-117 motif on GSK3β ^55^. Several studies identified a role for FRAT1 in the nuclear export of GSK3β ^33,56,57^. Hence, it is possible that the mTORC1-dependent and mTORC1-independent, GATOR1-dependent mechanisms regulate distinct aspects of GSK3β nuclear import or export, which will be of interest to examine in future studies.

How could GATOR1 and GATOR2 transmit signals to regulate GSK3β nucleocytoplasmic shuttling independently of mTORC1? Recent studies showed that the RING domains of certain GATOR2 complex proteins are essential for integrity of the GATOR2 complex ^17,18^. Moreover, the RING-domain protein WDR24 within GATOR2 supports ubiquitinylation of the GATOR1 subunit NPRL2 in leucine-rich conditions, leading to mTORC1 activation ^18^. It is intriguing to speculate that the GATOR2-dependent ubiquitinylation of NPRL2 upon leucine addition could also contribute to the TORC1-independent mechanism of regulation of GSK3β nuclear localization, although this has yet to be examined and should be a priority for future studies.

### Nuclear translocation of GSK3β supports cell survival in amino acid scarcity

Amino acid starvation resulting from the incubation of cells in EBSS media lacking serum yet supplemented with 10 ng/mL EGF showed a robust suppression in cell proliferation compared to cells grown in full media supplemented with serum (**Figure 7C-D**). This growth suppressive effect was observed irrespective of the localization status of GSK3β, as it was observed in the MDA-MB-231 WT cells, as well as MDA-MB-231 GSK3β-NES and GSK3β-NLS cells. In wild-type MDA-MB-231 cells, this loss of proliferation was not associated with changes to cell viability. This is consistent with engagement of cellular adaptive mechanisms that allow cell survival while arresting the cell cycle in the absence of amino acids from the extracellular environment.

The most well-understood adaptive mechanisms to amino acid scarcity result from inactivation of mTORC1 under these conditions, which can trigger a range of outcomes on cell physiology. This includes induction of autophagy that can replenish a source of amino acids derived from the hydrolysis of existing proteins (reviewed by ^58^). Our results show that control of GSK3β nucleocytoplasmic shutting also contributes to cell survival during amino acid starvation. Specifically, the translocation of GSK3β into the nucleus upon amino acid starvation (**Figure 1**), which is associated with a loss of GSK3β from the lysosomal membrane (**Figure 2**) may be required to sustain cell viability during amino acid starvation. This may result from the ability of nuclear GSK3β to phosphorylate nuclear substrates, expected upon amino acid starvation ^34^. Specifically, nuclear GSK3β can phosphorylate c-myc on T58, leading to c-myc degradation ^34^. The loss of c-myc in cancer cells leads to loss of cell proliferation and can trigger differentiation or loss of tumorigenicity ^59^, suggesting that nuclear GSK3β may regulate transcription to promote survival at the expense of proliferation.

Nuclear exclusion of GSK3β enhances sensitivity of cancer cells to rapamycin treatment ^33^ and transcription of genes involved in serine biosynthesis and one-carbon cycle while rendering cancer cells more sensitive to inhibition of SHIN1, a critical one-carbon metabolism enzyme ^39^. These studies, taken together with our results, suggest that upon amino acid starvation the nuclear translocation of GSK3β may suppress serine metabolism. However, amino acid starvation increases expression of enzymes involved in serine metabolism to promote cell survival and tumor progression ^60^. This suggests that the many other genes regulated by nuclear GSK3β ^39^ may promote survival specifically under amino acid starvation conditions. While the actions of GSK3β in the nucleus may thus be complex, targeting of GSK3β to the nucleus may induce a collective set of transcriptional programs that overall suppress proliferation and growth but enhance cell survival under periods of amino acid deprivation.

It may also be possible the loss of cytosolic GSK3β, rather than the gain in nuclear GSK3β under conditions of amino acid starvation, promotes cell survival. Several studies report that GSK3β promotes apoptosis (reviewed by ^61,62^), for example by phosphorylation of Bax ^63^, suggesting that nuclear accumulation of GSK3β and the resulting cytosolic depletion of GSK3β could suppress apoptosis under amino acid scarcity. Several studies suggested that GSK3β also suppresses autophagy ^61,64–66^, which may be related to regulation of endocytic membrane traffic and lysosomal acidification by GSK3β ^64,67^. These studies highlight that loss of GSK3β from the cytoplasm or specifically from the lysosome upon amino acid starvation may be required to enhance cell survival upon amino acid starvation.

In summary, we find that amino acid sensing that converges on GATOR1 and GATOR2 exerts control over GSK3β nucleocytoplasmic shutting. Surprisingly, this amino acid-dependent regulation of GSK3β nucleocytoplasmic shutting is mTORC1-independent, suggesting that GATOR1 signaling bifurcates from control of mTORC1 to regulate GSK3β. By experimentally restricting GSK3β to either the nucleus or cytosol, we also find that regulation of GSK3β nucleocytoplasmic shuttling by nutrient signaling may regulate cell survival. This work reveals new insight into how an mTORC1-independent amino acid sensing pathway modulates cell physiology.

## Materials and Methods

### Materials

Materials used in this study are described within each experimental method.

### Cell Culture

Wild-type human retinal pigment epithelial (ARPE-19) cells were cultured in Dulbecco’s Modified Eagle’s Medium/F-12 (DMEM/F12; Thermo Fisher Scientific) supplemented with 10% fetal bovine serum (Thermo Fisher Scientific) and 100 U/mL penicillin and 100 μg/mg streptomycin (Thermo Fisher Scientific). MDA-MB-231 cells were cultured in Dulbecco’s Modified Eagle’s Medium (ThermoFisher Scientific) supplemented with 10% fetal bovine serum (ThermoFisher Scientific) and 100 U/mL penicillin and 100 μg/mg streptomycin (ThermoFisher Scientific). Cells were incubated at 37℃ and 5% CO_2_ and were passaged upon reaching 80% confluency. For passaging, cells were washed with sterile phosphate-buffered saline (Sigma Aldrich, Oakville, ON) and lifted with 0.25% Trypsin-EDTA (Thermo Fisher Scientific).

### Stable transfections using Sleeping Beauty transposon system

pSBtet-BP was a gift from Eric Kowarz (Addgene plasmid # 60496 ; http://n2t.net/addgene:60496 ; RRID:Addgene_60496) ^68^. pCMV(CAT)T7-SB100 was a gift from Zsuzsanna Izsvak (Addgene plasmid # 34879 ; http://n2t.net/addgene:34879 ; RRID:Addgene_34879) ^69^. An oligonucleotide encoding DEPDC5 fused to eGFP was generated by BioBasic Inc (Markham, ON, Canada), using the sequence of DEPDC5 (obtained from vector V10849), starting with ATG AGA ACA ACA AAG GTC and ending with CAT GCC AGT GCC CCG) followed by the sequence encoding a spacer peptide (TCC GGA CTC AGA TCT CGA GCT CAA GCT), followed by the sequence encoding eGFP. This oligonucleotide sequence was subcloned into pSB-tet-BP to generate pSB-tet-BP-DEPDC5-BP. Oligonucleotides encoding GSK3b fused to an NLS or NES were generated as follows: the GSK3b sequence from NM_002093 (staring with: ATG TCA GGG CGG CCC AGA and ending with: TCA GCT TCC AAC TCC ACC), was fused to a segment encoding a spacer peptide and an HA tag (TCC GGA CTC AGA TCT CGA GCT CAA GCT TAC CCG TAT GAT GTT CCG GAT TAC GCT GGC TAT CCC TAC GAC GTG CCC GAC TAT GCC GGG TAC CCC TAT GAC GTC CCA GAC TAC GCA GCT), followed by either a NLS (CCT GCT GCG AAG CGC GTA AAA CTG GAC) as in ^70^, or an NES (CTG CAA AAA AAG TTG GAA GAG CTG GAA CTG) as in ^71^.

pSBtet-BP plasmids encoding various AAK1 WT and AAK1 mutant constructs alongside pCMV(CAT)T7-SB100 were co-transfected into ARPE-19 cells using FuGene HD transfection reagent (Promega), as per manufacturer’s protocol (Promega, Madison, WI), followed by a selection of stably engineered cells in media supplemented with 2 µg/mL puromycin for a period of 2-3 weeks.

pSBtet-BP stable cells were kept in cell culture media as indicated above but were incubated with 10% fetal bovine serum without tetracycline and maintained in 2 µg/mL puromycin. For the induction of DEPDC5-eGFP, Doxycycline (Dox) was used at a final concentration of 250 ng/mL in the cell culture media for 24 hours before downstream applications. For the induction of GSK3β-NES or GSK3β-NLS, Doxycycline (Dox) was used at a final concentration of 250 ng/mL in the cell culture media for 24 hours before downstream applications.

### siRNA transfection

ARPE-19 cells were subject to siRNA gene silencing for a target (DEPDC5, NRPL2, Mios, RagA, RagB) and non-targeting control. The following siRNAs were obtained as pre-validated sequences from Horizon Discovery (ON-TARGETplus, Human): DEPDC5 (9681): J-020708-20; NPRL2 (10641): D-015645-17-0010; Mios (54468): D-021181-03. Custom siRNAs were synthesized with sense strand sequences to target specific genes as follows: control (non-targeting): CGU ACU GCU UGC GAU ACG G; Targeting RagA: GAG AUG AAC CUC AGG AAU U; targeting RagB: GUA UUG AAC CUG UGG GAU U.

siRNA gene silencing was performed using Lipofectamine RNAiMAX (ThermoFisher Scientific), as per the manufacturer’s instructions and as previously described ^72–74^. Each siRNA construct was transfected at 220 pmol/l precomplexed and incubated in the transfection reagent in Opti-MEM (Gibco, ThermoFisher Scientific) for 4 h. Subsequently, cells were washed and incubated in regular growth medium. siRNA transfections were performed twice (72 h and 48 h) prior to experiments being performed.

### Amino acid starvation treatment conditions

Unless otherwise indicated, treatments with amino acid starvation and re-feeding were performed as follows: a minimum of 24h after seeding, cells were washed and then incubated for 90 min with (i) full media (DMEM/F12 for ARPE-19 cells or RPMI for MDA-MB-231 cells, each supplemented with 10% FBS), (ii) 90 min in EBSS (no serum, representing the “amino acid starvation” condition), or (iii) 60 min in EBSS followed by 30 min in EBSS with either 0.4 mM Leucine, 0.4 mM arginine, or 0.4 mM non-essential amino acids (NEAA, Source), representing the “refeeding” condition. For experiments also involving rapamycin treatment, cells were incubated with the respective media containing rapamycin for 30 min before all above treatments, for a total of 2h of treatment in rapamycin-containing media.

### Immunofluorescence staining, microscopy, and analysis

Wide-field epifluorescence (**Figures 1, 2, 3, 4, 5, 7B, S1, S2, S3**) was performed on fixed samples as previously described ^34^. Following treatments as indicated, cells were washed 2x in ice-cold PBS, and then subjected to methanol fixation. Samples were then blocked in 5% bovine serum albumin (BioShop), with GSK3β (Cell Signaling cat #9832) and/or LAMP1 (Cell Signaling cat # 9091) specific primary antibodies, followed by staining with fluorophore-conjugated secondaries and DAPI, and then mounting on glass slides in fluorescence mounting medium (DAKO, Carpinteria, CA). Antibodies for GSK3β were previously validated using this staining protocol ^34^. Imaging was performed on an Olympus IX83 Inverted Microscope with a 100x objective, coupled to a Hamamatsu ORCA-Flash4.0 digital camera (Olympus Canada, Richmond Hill, ON).

To determine nuclear levels of GSK3β (**Figures 1, 3, 4, 5, 6, S1**), a region of interest corresponding to the nucleus (identified by DAPI signal) was drawn manually using ImageJ ^75^. The mean fluorescence intensity of this region was measured, followed by subtraction of the mean fluorescence intensity of region of background (determined as a region of the image not corresponding to any cell). 30-40 cells were thus measured per experiment to determine the median nuclear GSK3β fluorescence intensity in each condition in each experiment. The Manders’ coefficient (**Figure 2**) was determined using Just Another Colocalization Plugin ^76^ within ImageJ ^75^, as previously described ^34^.

Live cell imaging to detect localization of DEPDC5-eGFP was done as follows: following induction of DEPDC5-eGFP with 250 ng/mL doxycycline for 24h, cells were incubated with media supplemented with a suspension of 3 nm silver nanoparticles prepared as in ^50^, allowing fluid phase update of particles and their accumulation in the lysosome; we previously demonstrated that this enhances the signal of green fluorescence at the lysosome without perturbing lysosomal function ^50^. This was followed by labeling with Lysotracker Deep Red (ThermoFisher Scientific) for 15 min. Following these labeling steps, cells were then treated as indicated: 90 min full media, 90 min EBSS, or 60 min EBSS followed by 30 min in EBSS supplemented with 0.4 mM leucine. Spinning disk confocal microscopy (**Figure 3**) was performed using Quorum (Guelph, ON, Canada) Diskovery combination total internal reflection fluorescence and spinning-disc confocal microscope, operating in spinning disc confocal mode, and as previously described ^50^. Single frames of each region of interest were obtained while cells were maintained at 37 C and 5% CO2. This instrument is comprised of a Leica DMi8 microscope equipped with a 63×/1.49 NA objective with a 1.8× camera relay (total magnification 108×). Imaging was done using 488 nm or 637 nm laser illumination and 527/30 and 700/75 emission filleters, and acquired using a iXon Ultra 897 EM-CCD (Spectral Applied Research, Andor). The Pearson’s coefficient (**Figure 3**) was determined using Just Another Colocalization Plugin ^76^ within ImageJ ^75^, as previously described ^77^.

### Western blotting

This was performed as previously described ^78^. After treatments as indicated, cells were washed with ice-cold PBS. Then, whole cell lysates were prepared in 2X-Laemelli Sample buffer (0.5M Tris pH 6.8, Glycerol, 10% SDS) supplemented with 1 mM sodium orthovanadate, 10 mM okadaic acid and 20 mM of protease inhibitor. This was followed by syringing of lysates at least 5 times with a 27.5-gauge syringe, supplementation of lysates with 10% beta-mercaptoethanol (to ensure reducing conditions) and 5% bromophenol blue, and heating at 65°C for 15 minutes. Whole cell lysates were resolved by SDS-PAGE and then transferred to 0.2 m pore PVDF membrane (Immobilon, Millipore). The membrane was blocked with blot wash buffer containing 3% BSA, incubated with primary antibodies in 1% BSA in blot wash buffer at 4°C overnight, and then incubated with secondary HRP-conjugated antibody in 1% BSA in blot wash; extensive washing in blot wash buffer was performed between each of these stages. Bands were visualized with Luminata ECL chemiluminescence substrate (Millipore Sigma).

### Cell proliferation and survival assay

Cell abundance and non-viable cells were measured using an Incucyte SX5 environmentally controlled microscope as previously described ^52^. Following induction of construct expression with doxycycline for 24h, MDA-MB-231 cells were placed in full media or EBSS supplemented with 10 ng/mL EGF, and also further supplemented with CellTox CellTox Green Cytotoxicity Assay reagent (Promega G8741) by adding 1µL of the dye in 10mL of media. Cytotoxicity and viability were measured using an Incucyte SX5 Live-Cell Analysis System (Sartorius). Images were acquired at regular intervals under 10x magnification using brightfield phase contrast and 100ms exposure of 460nm excitation, 524nm emission for green fluorescence. Analysis was performed with Incucyte Plategraph for CellTox positive cells and phase confluence.

### Statistical Analysis

For data with multiple treatments and comparisons involved normalized data, statistical analysis was performed using a Kruskal-Wallis test with Dunn’s multiple comparisons test (**Figures 1A**). For data with two dependent variables, statistical analysis was performed using two-way ANOVA without Geisser-Greenhouse correction, using Holm-Šídák post-test for pairwise comparisons between conditions (**Figures 2, 4, 5, 6, 7D**). For data with only two conditions, statistical analysis was performed by Student’s t-test (**Figure S4A**). For data with multiple treatments (and that does not involve a condition serving to normalize the rest of the data), statistical analysis was performed by a one-way ANOVA with a Tukey post-test (**Figure S4B**).

## Acknowledgements.

This study was supported by Operating Grants from the Cancer Research Society to C.N.A. and A.C.G. This work was also supported by Natural Sciences and Engineering Research Council Discovery Grants to C.N.A. (RGPIN-2016-04371), S.I. (RGPIN-2018-04161) and R.J.B. (RGPIN-2020-04343) as well as Canada Foundation for Innovation JELF awards to R.J.B. and C.N.A. (32957 and 38151) and a Tier 2 Canada Research Chair to R.J.B. (950-232333). This work was also supported by funding from the Faculty of Science at Toronto Metropolitan University.

