## Supplemental Figures for "Control of GSK3β nuclear localization by amino acid signaling requires GATOR1 but is mTORC1-independent"

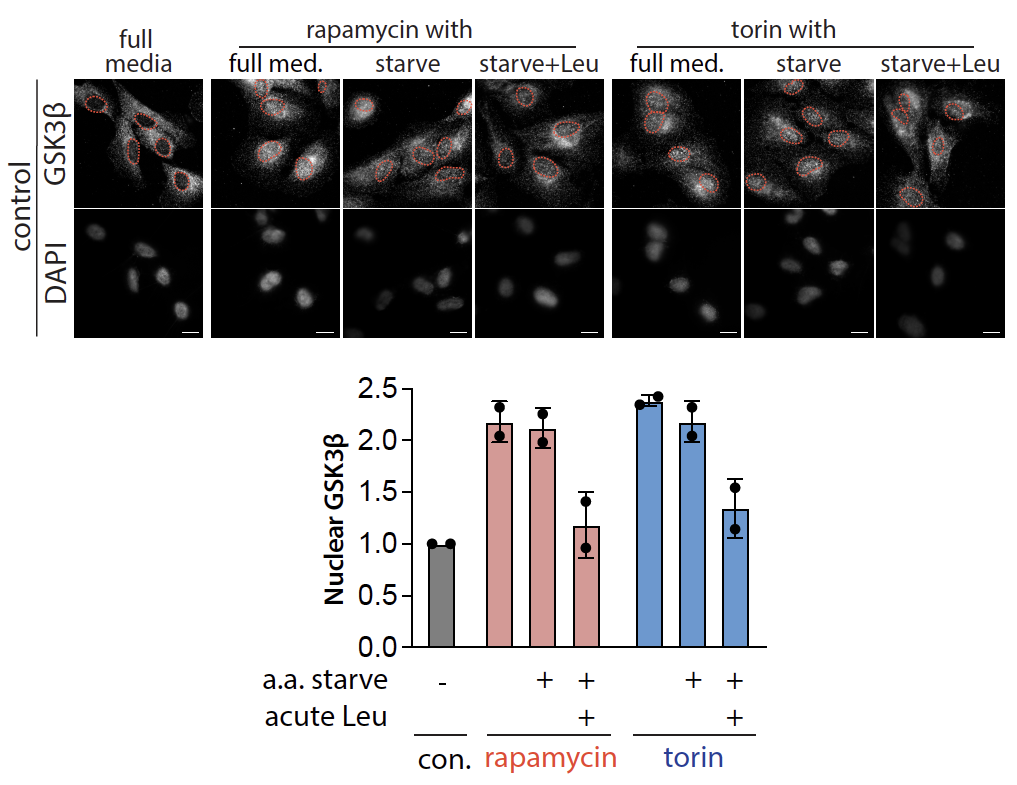

**Figure S1. Leucine stimulates GSK3β nuclear exit in starved ARPE-19 cells in the presence of rapamycin or Torin**. ARPE-19 cells were subjected to full media, amino acid starvation, or amino acid starvation followed by refeeding with leucine, with some samples also being subject to treatment with 1 µM rapamycin. Cells were fixed and labelled to detect endogenous GSK3β; shown (left panels) are representative micrographs obtained by widefield epifluorescence microscopy, scale 20 µm and (right panels) quantification of the mean fluorescence intensity of GSK3β in the nucleus depicting the mean ± SD from n = 2 independent experiments; each condition in each experiment determined by the median value from 30-40 individual cells.

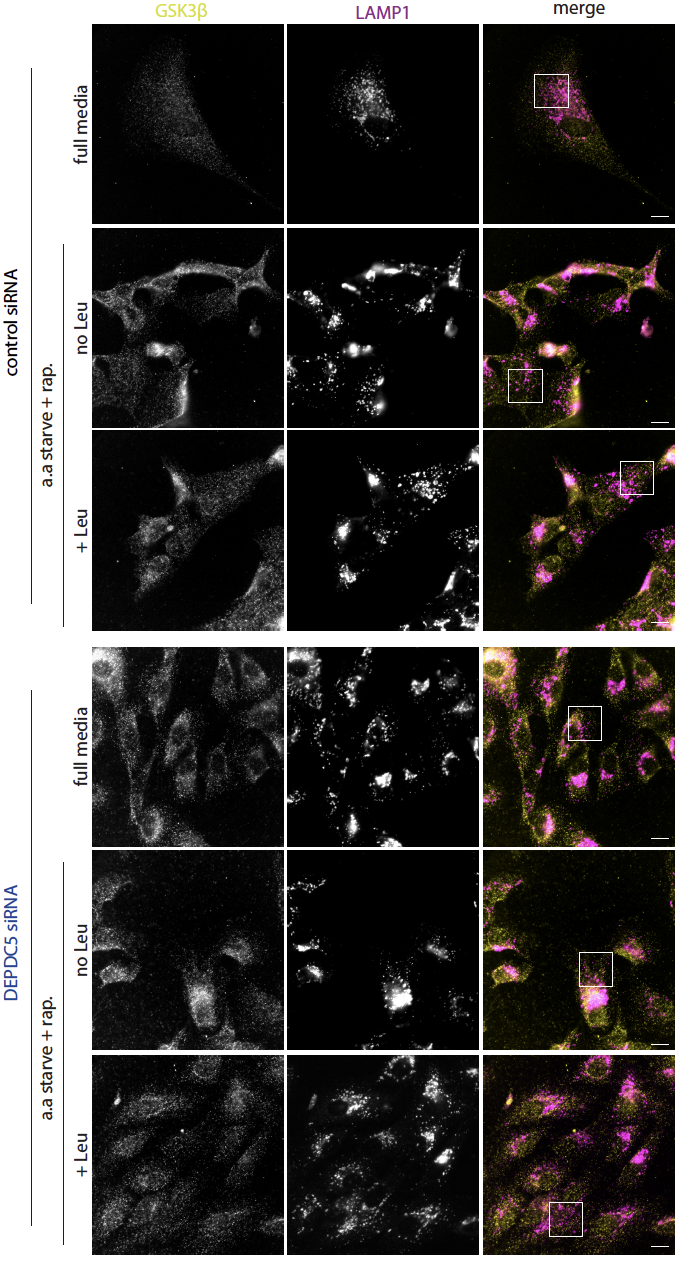

**Figure S2. Full image panels for Figure 2.** Shown are full-chip images corresponding to the images shown in Figure 2. Scale: 20 µm, white box: image region of interest shown in Figure 2.

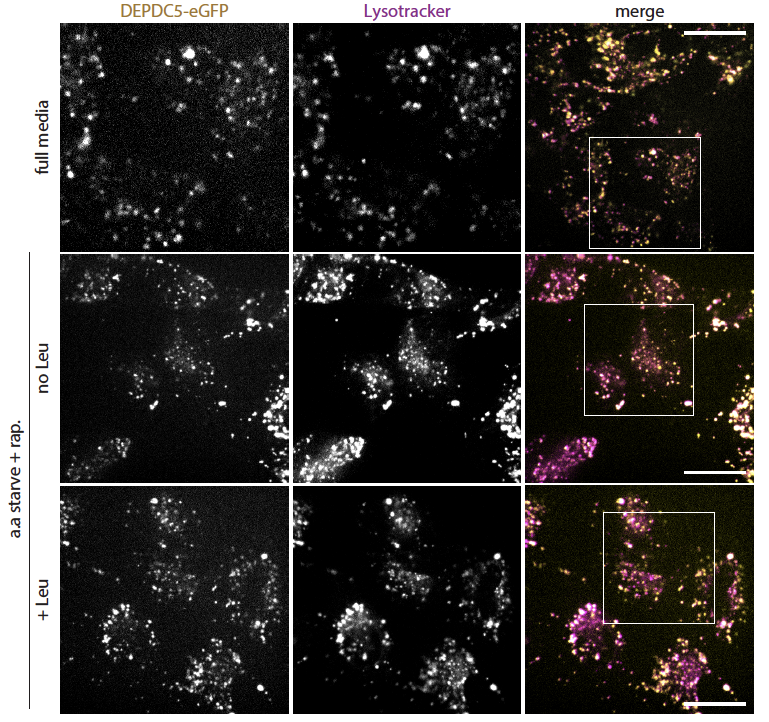

**Figure S3. Full image panels for Figure 3**. Shown are full-chip images corresponding to the images shown in Figure 3. Scale: 20 µm, white box: image region of interest shown in Figure 3.

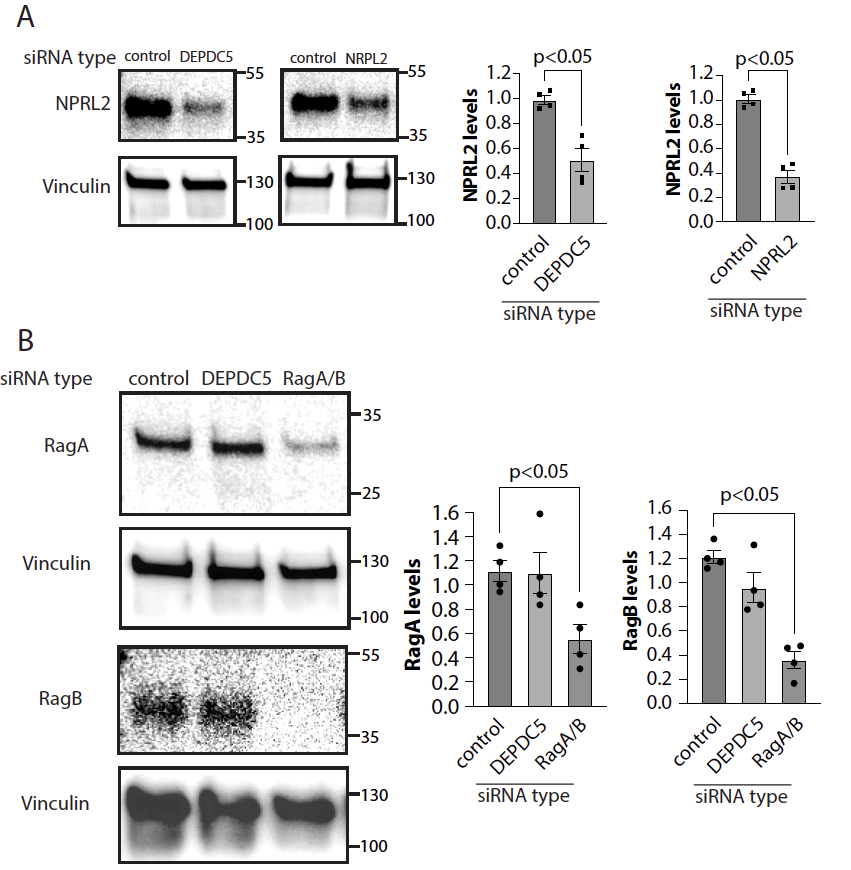

**Figure S4. Silencing of DEPDC5, NPRL2, and RagA/B in ARPE-19 cells**. ARPE-19 cells were transfected with siRNAs targeting both RagA and RagB, NPRL2, DEPDC5 or non-targeting siRNA, and then subjected to western blotting using antibodies as indicated. Levels of NPRL2 (A) and Rag A and Rab B (B) were quantified as described in Methods and shown relative to loading control (vinculin) as mean ± SEM from 4 independent experiments. Also shown for all western blot panels is the approximate molecular weight in kDa.
